# Development of a qPCR assay and a LAMP assay for *Verticillium longisporum* detection and a triplex qPCR assay for simultaneous detection of *V. longisporum*, *Leptosphaeria biglobosa* and *L. maculans* from canola samples

**DOI:** 10.1101/2024.01.24.577072

**Authors:** Heting Fu, Yalong Yang, Junye Jiang, Greg C. Daniels, Blake Hill, Shiming Xue, Kher Zahr, L. Stellar, Michael W. Harding, David Feindel, Carol Bvindi, Dilantha Fernando, Lipu Wang, Jie Feng

## Abstract

Verticillium wilt, Verticillium stem striping, and Verticillium stripe, are common disease names that all denote infection caused by *Verticillium longisporum*, on canola, or other Brassica crops. In this study, a quantitative PCR (qPCR) assay and a loop-mediated isothermal amplification (LAMP) assay were developed for the detection of *V. longisporum* from canola stem samples. Both assays are specific to *V. longisporum* at the species level and ubiquitous at the strain level. The low limit for positive detection of the two assays is 1 pg fungal DNA in a 20-µ L reaction or 1,400 fungal cells in 100-mg plant tissue. The qPCR assay was combined with the duplex qPCR assay for the two blackleg pathogens, *Leptosphaeria biglobosa* and *L. maculans* to constitute a triplex qPCR system for simultaneous detection of all three pathogens. The usefulness of this triplex qPCR system was verified on canola samples collected from various locations in Alberta, Canada. Using this triplex qPCR system, *V. longisporum* was detected from one sample, while the two blackleg pathogens were detected at higher frequencies. Since it is sometimes difficult to differentiate Verticillium stripe and blackleg on Alberta canola samples based on visual symptoms, the triplex qPCR system is an important tool for the detection of *V. longisporum*, especially when its presence is masked or obscured by symptoms of blackleg.

## Introduction

*Verticillium longisporum* is a serious pathogen on brassicaceous plants (Depotter et al. 2016). It has been reported in Europe (Karapapa et al. 1997), Russia (Pantou et al. 2005), Japan (Ikeda et al. 2012), China (Yu et al. 2015), and Canada (CFIA 2016). On oilseed rape (canola), the disease caused by *V. longisporum* has been called Verticillium wilt but later renamed to Verticillium stem striping (Depotter et al. 2016). For consistency and brevity, we will use the name “Verticillium stripe” throughout the remainder of the paper understanding that any of the aforementioned common disease names may apply to infections by this fungus.

In Canada, *V. longisporum* was first found in Manitoba in 2014, and a nationwide survey conducted by the Canadian Food Inspection Agency and the Canola Council of Canada indicated the presence of the pathogen in canola stubble from five other provinces: British Columbia, Alberta, Saskatchewan, Ontario and Quebec (CFIA 2016). However, thereafter *V. longisporum* on canola has not been confirmed in Alberta in annual canola disease surveys. Inspectors observed symptoms reminiscent of Verticillium stripe but, blackleg pathogens, *Leptosphaeria maculans,* and *L. biglobosa*, rather than *V. longisporum*, were isolated from the suspect plant samples (Harding et al. 2022; 2023).

Different from many other fungal species, *V. longisporum* is amphidiploid that resulted from hybridization between two of the following four haploid ancestors: genotype A1 of an unknown species, genotype D1 of another unknown species, and genotypes D2 and D3 of *V. dahliae*. The currently identified *V. longisporum* isolates can be classified into three lineages based on the parental combination: A1/D1, A1/D2, and A1/D3. Among the three lineages, A1/D1 is the most pathogenic lineage on canola, whereas lineage A1/D3 isolates are generally not pathogenic on this crop (Novakazi et al. 2015; Tran et al. 2013). Lineage A1/D2 is known only from horseradish in Illinois (USA) (Eastburn and Chang 1994). In Canada, a recent study indicated that *V. longisporum* isolates from canola and radish were in the A1/D1 group (Zou et al. 2020).

An effective disease management program is dependent on timely and proper identification of the causal agent of the disease. However, due to the diploid nature of *V. longisporum* and the fact that at least one of the parental species (i.e. *V. dahliae*) is a commonly occurring and widely distributed plant pathogen, accurate detection of *V. longisporum* has been difficult. Indeed, a DNA-based method for *V. longisporum* identification from plant samples has never been published. A multiplex PCR protocol developed by Inderbitzin et al. (2013) could be used to identify and differentiate *V. longisporum* into the lineage level by using DNA from pure fungal cultures. However, the use of this protocol for diagnosis of Verticillium stripe from plant samples has not been reported so far.

Quantitative PCR (qPCR) is an efficient method for plant disease diagnosis. qPCR especially probe-based qPCR is highly specific and sensitive. Another technique, loop-mediated isothermal amplification (LAMP), has gained popularity because of its capability for pathogen detection under a constant temperature and simple instrument requirements (Notomi et al. 2000). In recent years, LAMP reaction master mix became available commercially, which further facilitated the use of LAMP in diagnostics. Thus, in this study, we developed and evaluated a probe-based qPCR assay and a LAMP assay for specific detection of *V. longisporum* from infected canola plant samples. The qPCR assay was combined with a previously developed duplex qPCR for the two blackleg pathogens (Fu et al. 2023) to constitute a triplex qPCR system, which was then tested for capacity to specifically detect the three pathogens from canola samples collected from various fields in Alberta.

## Materials and methods

### Chemicals and standard techniques

All chemicals and instruments were purchased from Fisher Scientific Canada (Ottawa, ON) unless otherwise specified. All primers, probes, and gBlock were synthesized by Integrated DNA Technologies (Coralville, IA). All DNA extraction was conducted using the DNeasy Plant Pro kits (Qiagen Canada, Toronto, ON) with a Qiacube (Qiagen Canada). The extracted DNA was dissolved in 50 µ L water. When needed, DNA concentration was measured with a Nanodrop 1000. The whole-genome sequenced *V. longisporum* strains were identified by searching “Verticillium longisporum” in the National Center for Biotechnology Information (NCBI) genome database (https://www.ncbi.nlm.nih.gov/datasets/genome/?taxon=100787). The average genome size of all the whole-genome sequenced strains was calculated. The molecular weight of the average genome was calculated using an online ssDNA Mass to Moles Convertor (https://nebiocalculator.neb.com).

### Multiplex PCR, qPCR, and LAMP reactions

Multiplex PCR was conducted in Promega Go Taq Master Mix or Thermo Scientific Phire Plant Direct PCR Master Mix with a T100 thermocycler (Bio-Rad Canada, Mississauga, ON). Each 20-µ L reaction contained 0.25 µM of each primer and 1 ng DNA from fungal cultures or 20 ng DNA from plants (infected or healthy). The PCR program followed Zou et al. (2020). SYBR Green-based qPCR and probe-based qPCR were conducted in SsoAdvanced universal SYBR green supermix (Bio-Rad Canada) and PrimeTime gene expression master mix (Integrated DNA Technologies), respectively, in a CFX96 touch real-time PCR detection system (Bio-Rad Canada). Each 20-µ L qPCR reaction contained 0.25 µ M of each primer, 0.18 µ M of each probe (probe-based qPCR only), and 2 µ L of each DNA template regardless of the concentrations. Each qPCR reaction was conducted with three technical replicates. In each 96-well PCR plate, three replicates of negative-control reactions were also included, in each of which 2 µ L of water was used as a template. The qPCR program consisted of an initial denaturation at 95°C for 3 min, followed by 40 cycles of denaturation at 95°C for 10 s and annealing/extension at 60°C for 30 s. All LAMP reactions were conducted in the WarmStart® Colorimetric LAMP Master Mix (NEB Canada, Whitby, ON). Each 20-µ L reaction contained 2 µ L of template. Quantities of other components in each reaction followed the NEB’s instructions for the master mix. All reactions were conducted in individual 200-µ L PCR tubes. The reaction program consisted of only one step in which the tubes were incubated at 65°C for 30 minutes. After the incubation, the reactions were checked visually and the results were recorded by photography.

### Selection of genes for primer designing

Candidate genes for qPCR and LAMP primer design were selected by the following approach. All proteins of *V. longisporum* were accessed by searching the NCBI database (https://www.ncbi.nlm.nih.gov) with the term ‘Verticillium longisporum’, which ended up with 95,405 protein entries. From these proteins, those consisting of 50-60 amino acids were retrieved, which ended up with 366 entries. After removing the 13 proteins with predicted functions and 222 partial proteins, 131 protein entries remained. Blastp was conducted with these 131 proteins as queries against NCBI protein database. After removing the proteins that produced hit(s) from non-*V. longisporum* species, 66 proteins remained. The genomic sequences (i.e. coding sequence and introns) were retrieved for these 66 proteins (hereafter referred to as 66 genes). Two Blastn analyses were conducted using the genomic sequences of these 66 genes as queries, one against the nucleotide collection (nr/nt) database and the other against the whole-genome shotgun contigs (wgs) database with the Organism set as Fungi (taxid:4751). After removing the genes that produced hit(s) from non-*V. longisporum* species, four genes remained (GenBank accession numbers CRK29573, CRK26624, CRK12486, and CRK45493). Among these four genes, the first two are present in all the eight whole-genome sequenced *V. longisporum* strains (strains PD589, VLB2, VL20, VL2, Vl32, VL1, Vl43, and Vl145c) with 100% query cover and identical sequences. The other two were present in six whole-genome sequenced strains with 100% query cover and identical sequences but were not present in strains Vl32 and PD589. Both CRK29573 and CRK26624 had a coding sequence of 180 nucleotides (nt), with the former having no intron and the latter having a 58-nt intron (thus the gene is 238 nt).

### Primer, probe, and gBlock design for qPCR and LAMP

Based on the sequence of CRK29573, qPCR primer pairs were designed using Primer-Blast (https://www.ncbi.nlm.nih.gov/tools/primer-blast). Each of the designed primer pairs was tested in Primer 3 (https://primer3.ut.ee) for probe designing. Three primers/probe sets were obtained. The sequences of the primers and the probe of each of these three sets and the sequences of the primers/probe sets (P-Lm and P-Lb; Table 1) used for the duplex qPCR for canola blackleg detection (Fu et al. 2023) were analyzed for potential primer dimers using Multiple Primer Analyzer hosted by Thermo Fisher (https://www.thermofisher.com). One set without any predicted primer dimer was selected to be used in this study and named P-VL (Table 1). The probe in P-VL was labeled with the fluorescent dye Cyanine5 (Cy5). A DNA sequence at 499 nt, corresponding to nt 2669667 to 2670165 of the *V. longisporum* strain PD589 whole genome sequence (GenBank accession number 200JAETXT010000011) and containing the sequence of CRK29573, was retrieved from the NCBI database. A double-stranded DNA fragment of this 499-nt sequence was synthesized as gBlock and named G-VL.

**Table 1.**
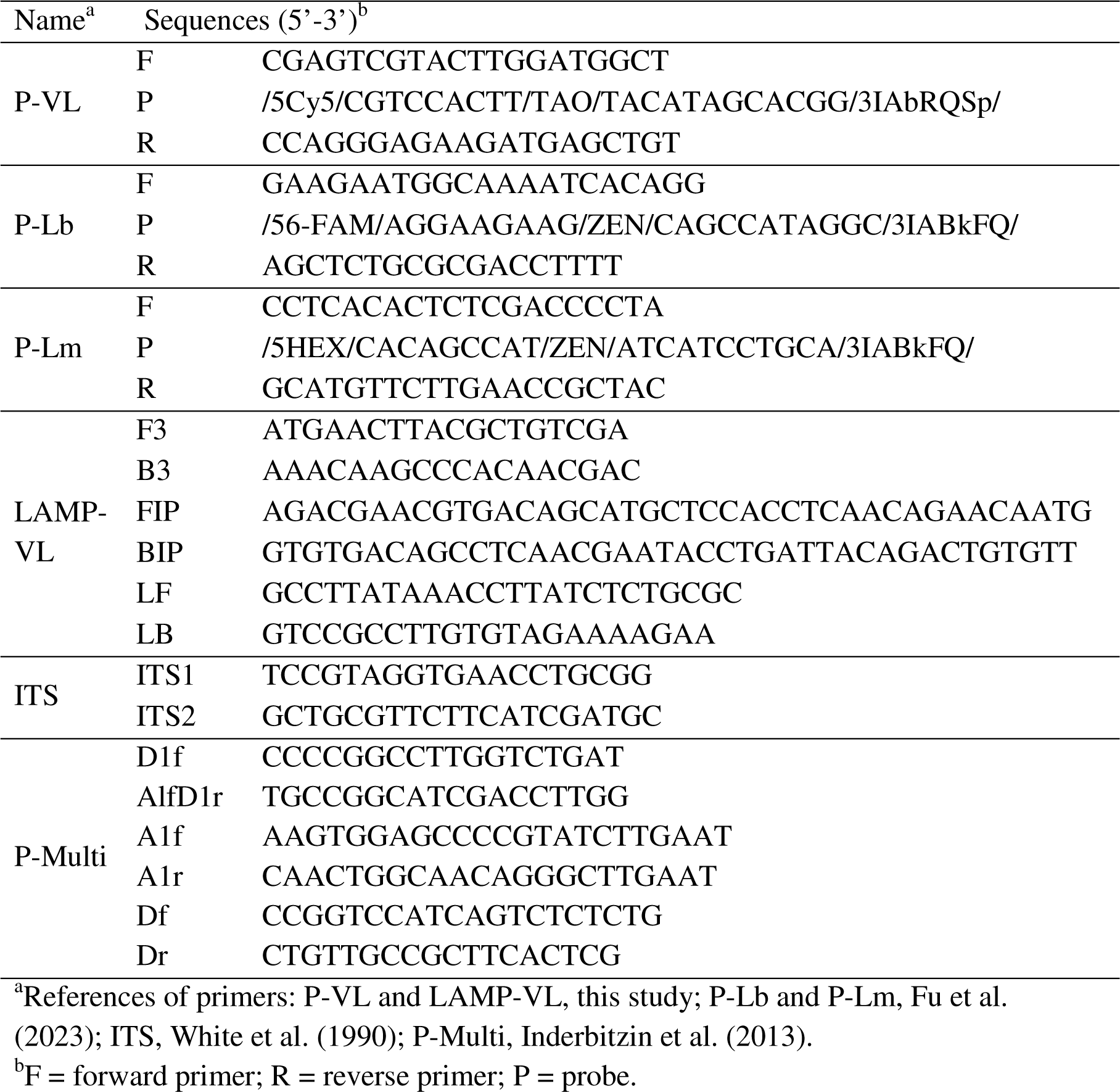
Primers and probes used in this study Namea Sequences (5’-3’)b.

Based on the sequence of CRK26624, two sets of LAMP primers were designed using Primer Explorer (https://primerexplorer.jp). After testing on *V. longisporum* genomic DNA, only one set was used in further studies and named LAMP-VL (Table 1).

### Specificity test of P-VL and LAMP-VL

The specificities of P-VL and LAMP-VL were tested on one known *V. longisporum* isolate (APHL 189), and 27 isolates of other species including 18 fungi and one oomycete (*Pythium intermedium*). APHL 189 was isolated from a canola sample provided by Dr. Manika Pradhan (Manitoba Agriculture, Food and Rural Development) in 2016. Other species are common pathogens of canola or have been isolated by Alberta Plant Health Lab (APHL) from either diseased canola samples or samples of crops rotated with canola in Alberta. All fungal and oomycete species were grown on 4% (w/v) potato dextrose agar (PDA) plates. For each species, approximately 50 mg of fresh mycelia was harvested and subjected to DNA extraction. DNA was also extracted from a canola clubroot gall sample infected by *Plasmodiophora brassicae* pathotype 5. In addition, four DNA samples were prepared as negative controls, including one from 7-day-old healthy canola leaves and one from 7-day-old healthy canola roots. The third and fourth negative controls were prepared from a pellet from 50 mL of YPG media (1% yeast extract, 1% peptone, 2% glucose, w/v) and LB broth, respectively, after incubating 10 g of canola root tissue collected from a *Verticillium*-free field plot at 150 rpm 30°C for 48 hours (the pellet was prepared by centrifuging the culture after the root tissue being removed at 4,000 × *g* for 5 min). The concentration of all DNA samples was adjusted to 500 pg/µ L. The qPCR primers/probe set P-VL and LAMP primer set LAMP-VL were tested on all the above-mentioned DNA samples. In addition, to have a DNA quality and quantity control, all DNA samples were tested with the universal primer pair ITS1/ITS2 (White et al. 1990; Table 1) by SYBR green-based qPCR. The qPCR analysis was conducted twice.

### Sensitivity test of qPCR and LAMP

The gBlock G-VL was dissolved in water and a 10 pM stock solution was prepared, which roughly equals 6 × 10^6^ double-stranded DNA copies/µ L. A set of 10-fold serial dilutions was prepared from 6 × 10^6^ copies/µ L to 6 copies/µ L. The serial dilutions were tested by probe-based singleplex qPCR using P-VL. The experiment was conducted twice with alternative preparations of gBlock serial dilutions.

The sensitivity of P-VL was further tested on genomic DNA of a known *V. longisporum* isolate APHL 189. Genomic DNA of APHL 189 was adjusted to 50 ng/µ L and then a set of 10-fold serial dilutions was prepared from 50 ng/µ L to 0.5 fg/µ L. The serial dilutions were tested by probe-based singleplex qPCR using P-VL and by triplex probe-based qPCR using P-VL, P-Lb, and P-Lm. P-Lb and P-Lm were developed by Fu et al. (2023) for the detection of the two canola blackleg pathogens *L. biglobosa* and *L. maculans*, respectively. The serial dilutions of the APHL 189 genomic DNA were also tested by LAMP using the primer set LAMP-VL. The qPCR and LAMP experiments were conducted twice with alternative preparations of genomic DNA serial dilutions.

### Test of qPCR and LAMP on field samples

Twenty-four canola stem samples showing black lesions were collected from 15 counties in central and southern Alberta (Fig. 1). In addition, two canola stem samples (MB 1 and MB 2) were collected from Manitoba, in which MB 2 was confirmed to be infected by *V. longisporum* (Dilantha Fernando, unpublished data). The samples were cut into 2-mm cubes and air-dried overnight at 23°C. From each of the air-dried samples, a 100-mg subsample was subjected to DNA extraction. The obtained DNA samples were tested by the triplex qPCR using P-VL, P-Lb and P-Lm and LAMP using LAMP-VL. The experiment was conducted twice on DNA samples from alternative 100-mg subsamples for each plant sample.

**Fig. 1.**
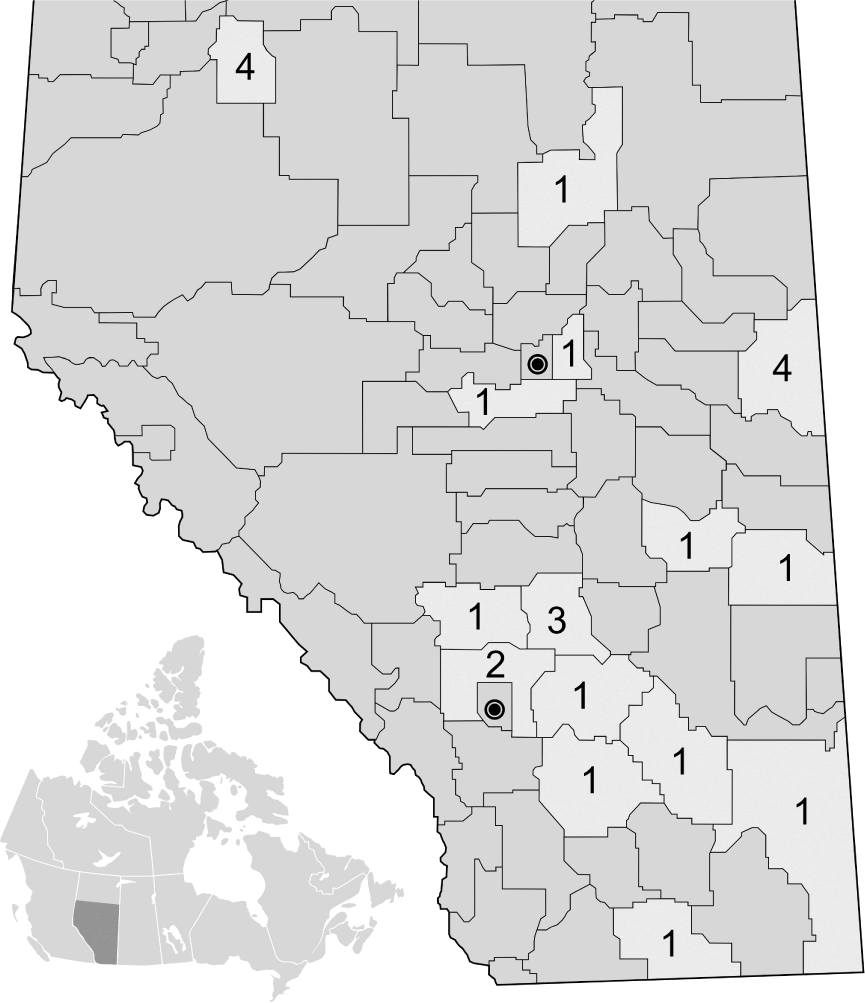
Geographic locations in Alberta counties of the 24 canola stem samples that were collected for Verticillium longisporum detection. In each county, the number of samples was indicated.

### Test of multiplex PCR on field samples and *V. longisporum* isolates

Selected field samples and the *V. longisporum* isolate APHL 189 were tested by the multiplex PCR using the primer set P-Multi (Table 1), which was developed by Inderbitzin et al. (2013) and included three primer pairs specific to the glyceraldehyde-3-phosphate dehydrogenase gene (GAPDH), the elongation factor 1-alpha gene (EF1) and the internal transcribed spacer region (ITS), respectively. According to Inderbitzin et al. (2013), from the *V. longisporum* lineage A1/D1, the EF1 and ITS primers will produce a 310-base pair (bp) band and a 1020-bp band, respectively; from the lineage A1/D2, the EF1 primers will produce the 310-bp band; from the lineage A1/D3, the GAPDH and EF1 primers will produce a 490-bp band and the 310-bp band, respectively and from *V. dahliae*, only the GAPDH primers will produce the 490-bp band. Thus, the primer set P-Multi can be used to differentiate the three *V. longisporum* lineages and *V. dahliae* from each other.

## Results

### Specificity of P-VL and LAMP-VL

In the qPCR analysis, the primer set ITS1/ITS2 produced signals from all the DNA samples, as indicated by the quantification cycle (Cq) value from each reaction (Table 2), which provided a reference for the quality and quantity of each DNA sample. P-VL produced signals from the *V. longisporum* isolate APHL 189 (Table 2). In contrast, no signal was produced from any of the triplicated qPCR reactions on DNA samples of all the non-*V. longisporum* isolates and the four negative controls (Table 2). In the LAMP analysis, LAMP-VL generated identical results as P-VL in the qPCR analysis: the signal was produced from the *V. longisporum* isolate APHL 189 only (data not shown). Similar results were obtained from the repeated experiment. These results confirmed that both P-VL and LAMP-VL are specific to *V. longisporum*.

**Table 2.** Specificity test of the qPCR primer set P-VL on canola pathogens.

| Species | Strain<sup>a</sup> | Cq values of qPCR<sup>b</sup> |  |
| --- | --- | --- | --- |
|  |  | ITS1/ITS2 | P-VL |
| *Alternaria infectoria* | APHL 002 | 24.2±0.1 | NS |
| *Alternaria solani* | APHL 004 | 21.0±0.2 | NS |
| *Botrytis cinerea* | APHL 199 | 19.0±0.1 | NS |
| *Colletotrichum coccodes* | APHL 029 | 28.2±0.3 | NS |
| *Fusarium oxysporum* | APHL 069 | 24.3±0.1 | NS |
| *Leptosphaeria biglobosa* | APHL 216 | 21.1±0.1 | NS |
| *Leptosphaeria biglobosa* | LB19-7 | 17.4±0.1 | NS |
| *Leptosphaeria biglobosa* | APHL 1170 | 17.1±0.1 | NS |
| *Leptosphaeria biglobosa* | LB19SK-11-2 | 18.0±0.1 | NS |
| *Leptosphaeria maculans* | LM19-11 | 18.2±0.2 | NS |
| *Leptosphaeria maculans* | LM19SK-11-21 | 18.3±0.1 | NS |
| *Leptosphaeria sclerotioides* | APHL 101 | 18.2±0.1 | NS |
| *Phoma aliena* | APHL 128 | 20.3±0.1 | NS |
| *Phoma betae* | APHL 130 | 20.1±0.1 | NS |
| *Phoma medicaginis* | APHL 134 | 19.4±0.1 | NS |
| *Plasmodiophora brassicae* | Pathotype 5 | 25.8±0.1 | NS |
| *Pythium intermedium* | APHL 159 | 18.0±0.2 | NS |
| *Rhizoctonia solani* | APHL 234 | 22.9±0.2 | NS |
| *Sclerotinia sclerotiorum* | APHL 244 | 19.7±0.1 | NS |
| *Stagonospora nodorum* | APHL 120 | 21.6±0.1 | NS |
| *Trichoderma sp.* | APHL 242 | 23.6±0.1 | NS |
| *Verticillium albo-atrum* | APHL 190 | 18.3±0.2 | NS |
| *Verticillium albo-atrum* | APHL 191 | 20.2±0.2 | NS |
| *Verticillium albo-atrum* | APHL 192 | 18.3±0.1 | NS |
| *Verticillium dahliae* | APHL 232 | 17.4±0.1 | NS |
| *Verticillium dahliae* | APHL 303 | 23.0±0.2 | NS |
| *Verticillium lecanii* | APHL 032 | 22.7±0.3 | NS |
| *Verticillium longisporum* | APHL 189 | 21.5±0.2 | 25.8±0.1 |
| *Verticillium tricorpus* | APHL 1117 | 22.8±0.1 | NS |
| Negative control | Canola leaf | 21.2±0.1 | NS |
| Negative control | Canola root in LB<sup>c</sup> | 24.2±0.1 | NS |
| Negative control | Canola root in YPG<sup>c</sup> | 26.6±0.1 | NS |
| Negative control | Canola root | 20.4±0.1 | NS |

<sup>a</sup>All strains belong to Alberta Plant Health Lab plant pathogen culture collection, a project funded by the Canadian Agricultural Partnership (no. 601322) to JF except: Lb19SK-11-2 and LM19SK-11-21 were provided by Dr. Gary Peng from Agriculture and

### Sensitivity of P-VL in singleplex qPCR

The serial dilutions of the gBlock G-VL and *V. longisporum* genomic DNA were tested in singleplex qPCR with P-VL. Based on the Cq values, standard curves were constructed for P-VL against gBlock (Fig. 2a) and genomic DNA (Fig. 2b). The efficiencies of P-VL against gBlock and genomic DNA were 1.01 (Fig. 2a) and 1.00 (Fig. 2b), respectively.

The low limit of P-VL for a positive detection in a 20-µ L reaction was 24 (10^1.08^ × 2 = 24) single-stranded gBlock copies (Fig. 2a) or 1 pg genomic DNA (Fig. 2b). The average genome size of the eight whole-genome sequenced *V. longisporum* stains is 69.7 Mb. Based on the molecular weight calculation, one pg of the genomic DNA is composed of 28 genomes of *V. longisporum*. Thus the low detection limits of the qPCR assay using P-VL were similar against gBlock and fungal genomic DNA.

**Fig. 2.**
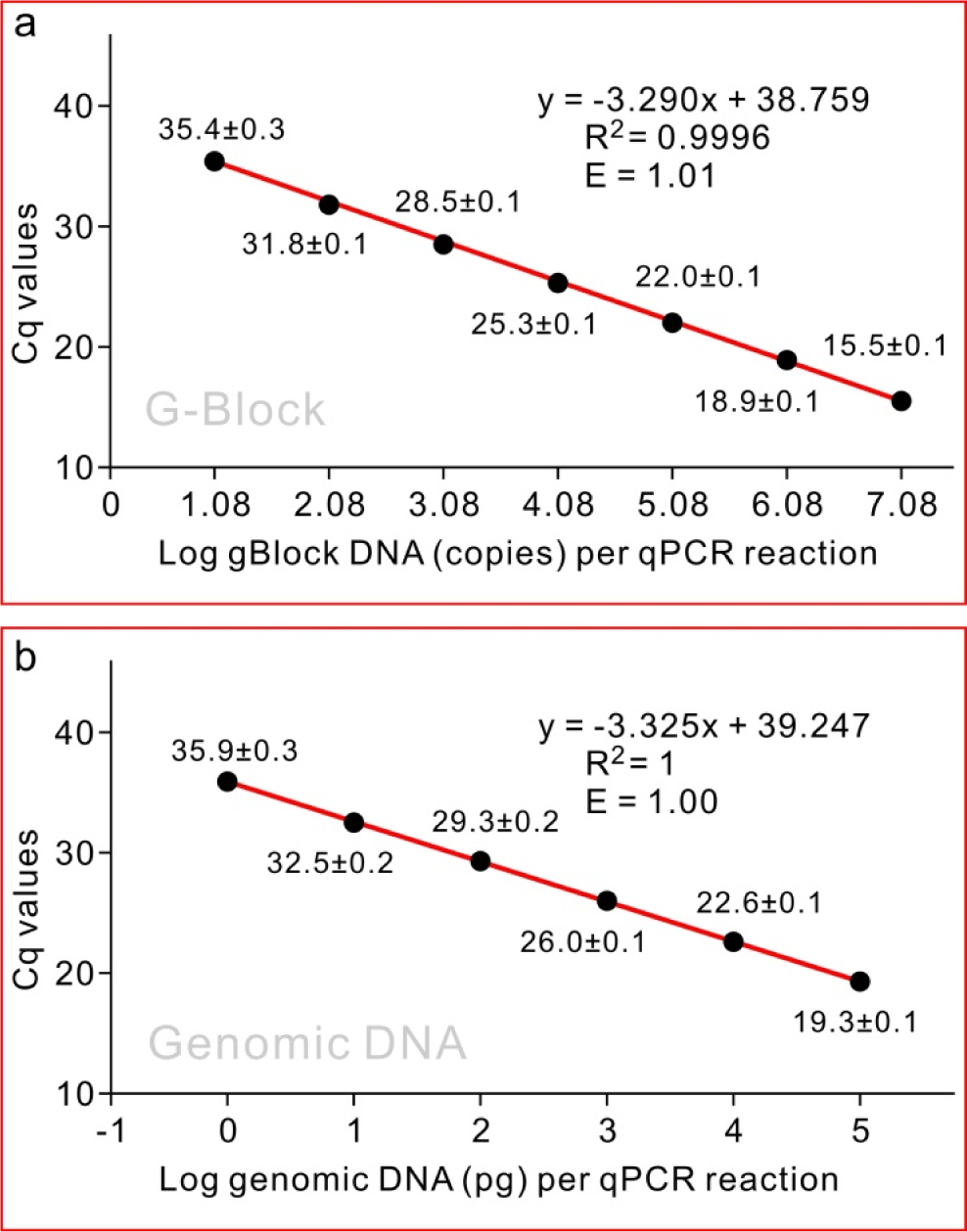
Sensitivity of the qPCR primer set P-VL. a, on gBlock DNA. b, on Verticillium longisporum genomic DNA. The qPCR standard curves were generated from the mean of quantification cycle (Cq) values against log10 of gBlock double-stranded DNA copies (a) or picogram of genomic DNA (b) in one reaction. The R2 score of the equation and the efficiency of the primers (E) are indicated over the curve. Efficiency was calculated as E = −1+10(−1/slope). Each data point is shown as mean of three technical replicates ± standard deviation.

### Sensitivity of LAMP-VL

The serial dilutions of the *V. longisporum* genomic DNA were tested by LAMP using LAMP-VL. The low limit of LAMP-VL for a positive detection in a 20-µ L reaction was 1 pg genomic DNA (Fig. 3). This result indicated that the LAMP system has a similar sensitivity as the qPCR system.

**Fig. 3.**
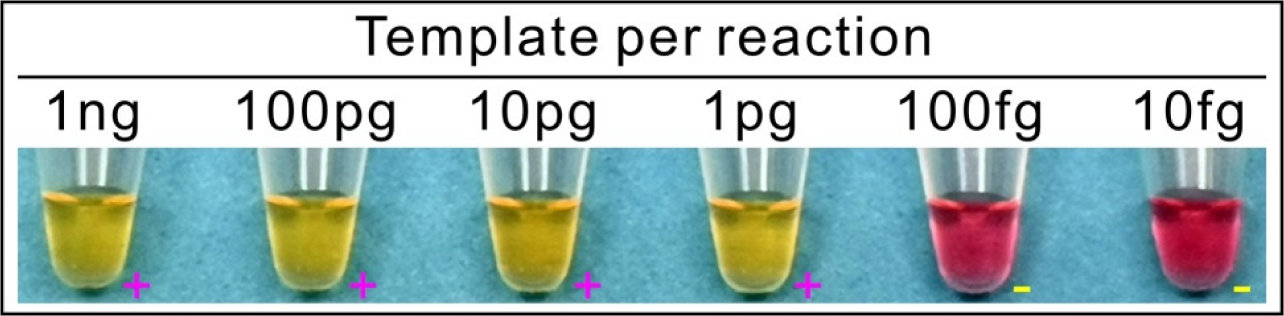
Sensitivity of the LAMP primer set LAMP-VL. The LAMP reactions were conducted against a set of serial dilutions of Verticillium longisporum genomic DNA, from 1 ng per reaction to 10 fg per reaction. Changing of the color of the reaction mixture from red to yellow indicated a positive reaction. Each tube was also indicated by a + for positive reaction or a – for negative reaction.

### Sensitivity of P-VL in triplex qPCR

The serial dilutions of the *V. longisporum* genomic DNA were tested in triplex qPCR using P-VL, P-Lb, and P-Lm. Based on the Cq values, standard curves were constructed for each primer set (Fig. 4a-c). The efficiencies of P-VL, P-Lb, and P-Lm were 1.02, 1.00 and 0.97, respectively. The low limits of genomic DNA for a positive detection in a 20-µ L reaction were 1 pg for P-VL (Fig. 4a), 0.4 (10^−0.42^ = 0.4) pg for P-Lb (Fig. 4b) and 0.5 (10^−0.27^ = 0.5) pg for P-Lm (Fig. 4c). This result indicated that the sensitivity of P-VL in triplex qPCR was similar to that in singleplex qPCR.

**Fig. 4.**
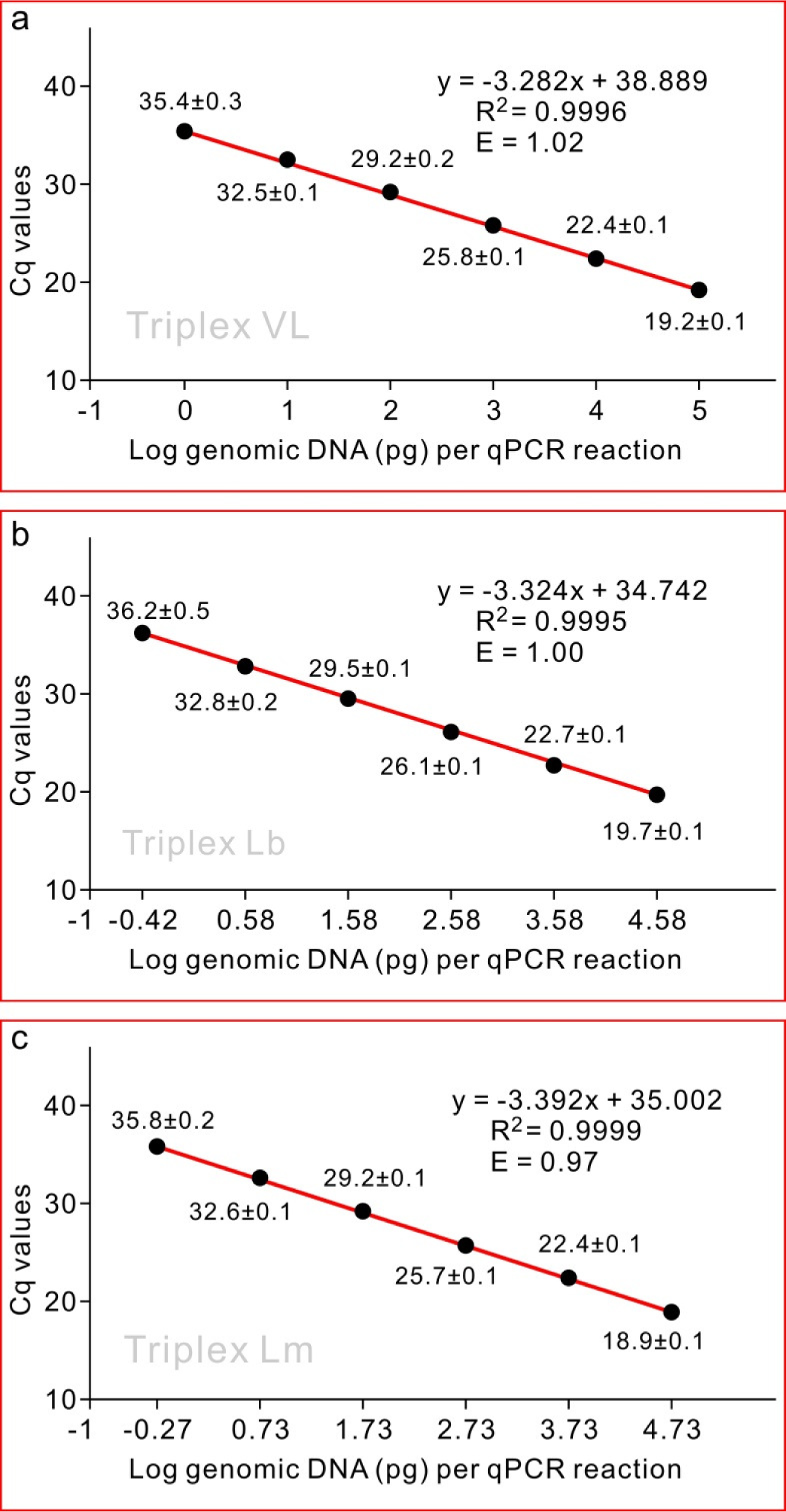
Sensitivity of the triplex qPCR system for the detection of Verticillium longisporum by the primer set P-VL (a), Leptosphaeria biglobosa by the primer set P-Lb (b) and L. maculans by the primer set P-Lm (c) in one reaction. The qPCR standard curves were generated from the mean of quantification cycle (Cq) values against log10 of picograms of serial dilutions of genomic DNA from each of the three fungi. The R2 score of the equation and the efficiency of the primers (E) are indicated over the curve. Efficiency was calculated as E = −1+10(−1/slope). Each data point is shown as mean of three technical replicates ± standard deviation.

### Triplex qPCR and LAMP analyses on field samples

Twenty-six canola samples, 24 from Alberta and two from Manitoba were tested by the triplex qPCR using the primer sets P-VL, P-Lb, and P-Lm (Table 3). *Verticillium longisporum* was detected from one Alberta sample (23CAN-305B) and one Manitoba sample (MB 2). Of the three pathogens, *V. longisporum*, *L. biglobosa,* and *L. maculans*, at least one was detected from each of the 26 samples. From 13 of the 24 Alberta samples, both *L. biglobosa* and *L. maculans* were detected. Among the 13 samples, the sample positive for *V. longisporum* (23CAN-305B) was also positive for *L. biglobosa* and *L. maculans*. In addition, all 26 canola samples were tested by LAMP using LAMP-VL. The LAMP results were identical to the qPCR results for the yes/no detection of *V. longisporum* (Table 3). These results indicated that both the triplex qPCR and LAMP developed in this study can be used on field samples.

**Table 3.** Triplex qPCR and LAMP analyses on canola samples.

| Samples | Location | Cq values of triplex qPCR<sup>a</sup> |  |  | LAMP<sup>b</sup> |
| --- | --- | --- | --- | --- | --- |
|  |  | P-Lb | P-Lm | P-VL |  |
| 2025A | Alberta | 25.1±0.1 | 24.7±0.1 | NS | - |
| 2025B | Alberta | 31.5±0.2 | 29.0±0.3 | NS | - |
| 2025C | Alberta | 32.3±0.2 | NS | NS | - |
| 2025D | Alberta | 26.5±0.1 | NS | NS | - |
| 23CAN-023 | Alberta | NS | 23.8±0.1 | NS | - |
| 23CAN-042 | Alberta | NS | 23.5±0.1 | NS | - |
| 23CAN-079 | Alberta | 28.0±0.1 | 33.2±0.4 | NS | - |
| 23CAN-083 | Alberta | 37.2±0.1 | 24.1±0.1 | NS | - |
| 23CAN-099 | Alberta | 28.3±0.1 | 25.1±0.1 | NS | - |
| 23CAN-135 | Alberta | 37.0±0.7 | NS | NS | - |
| 23CAN-152 | Alberta | 31.6±0.2 | NS | NS | - |
| 23CAN-197 | Alberta | 24.8±0.1 | 28.4±0.1 | NS | - |
| 23CAN-199 | Alberta | 27.0±0.1 | NS | NS | - |
| 23CAN-207 | Alberta | 30.3±0.1 | 38.5±0.9<sup>c</sup> | NS | - |
| 23CAN-207 | Alberta | 28.3±0.1 | 27.6±0.1 | NS | - |
| 23CAN-305B | Alberta | 26.9±0.1 | 26.7±0.1 | 30.0±0.1 | + |
| 23CAN-305C | Alberta | 29.7±0.1 | 23.0±0.1 | NS | - |
| 23CAN-310 | Alberta | NS | 25.5±0.1 | NS | - |
| 23CAN-321 | Alberta | 37.2±0.8<sup>c</sup> | 23.0±0.1 | NS | - |
| 23CAN-325 | Alberta | 24.2±0.1 | 37.2<sup>c</sup> | NS | - |
| 23CAN-333A | Alberta | NS | 25.3±0.1 | NS | - |
| 23CAN-342 | Alberta | 36.4±0.6 | NS | NS | - |
| 23CAN-348 | Alberta | 24.2±0.1 | NS | NS | - |
| 23CAN-363 | Alberta | 31.4±0.1 | 22.2±0 | NS | - |
| MB 1 | Manitoba | NS | 22.7±0.1 | NS | - |
| MB 2 | Manitoba | 24.1±0.1 | NS | 23.2±0.1 | + |
| Negative control<sup>d</sup> |  | NS | NS | NS | - |
| Water |  | NS | NS | NS | - |

<sup>a</sup>Triplex probe-based qPCR using the primer sets P-Lb, P-Lm and P-VL. Quantification cycle (Cq) data were shown as mean ± standard deviation (n = 3). NS = no signal from all the three replicates.
<sup>b</sup>LAMP analysis using the primer set LAMP-VL. +, positive reaction; -, negative reaction.
<sup>c</sup>Only one (without a standard deviation value) or two (with a standard deviation value) of the three technical replicates generated Cq values.
<sup>d</sup>DNA from root of a 14-day old healthy canola seedling.

### Confirmation of the triplex qPCR and LAMP results by multiplex PCR

With the Promega master mix, the multiplex PCR produced a single band close to 490 bp from the two *V. dahliae* isolates, two bands close to 1020 bp and 310 bp, respectively, from the *V. longisporum* isolate and three *V. longisporum*-infected canola samples (Fig. 5a). With the Thermo Fisher master mix, the multiplex PCR produced similar results as with the Promega master mix except that two bands were produced from the two *V. dahliae* isolates (Fig. 5b). Nevertheless, multiplex PCR confirmed the *V. longisporum* identity of APHL 189 and the presence of *V. longisporum* in the three canola samples, and also confirmed the specificity of our triplex qPCR and LAMP.

**Fig. 5.**
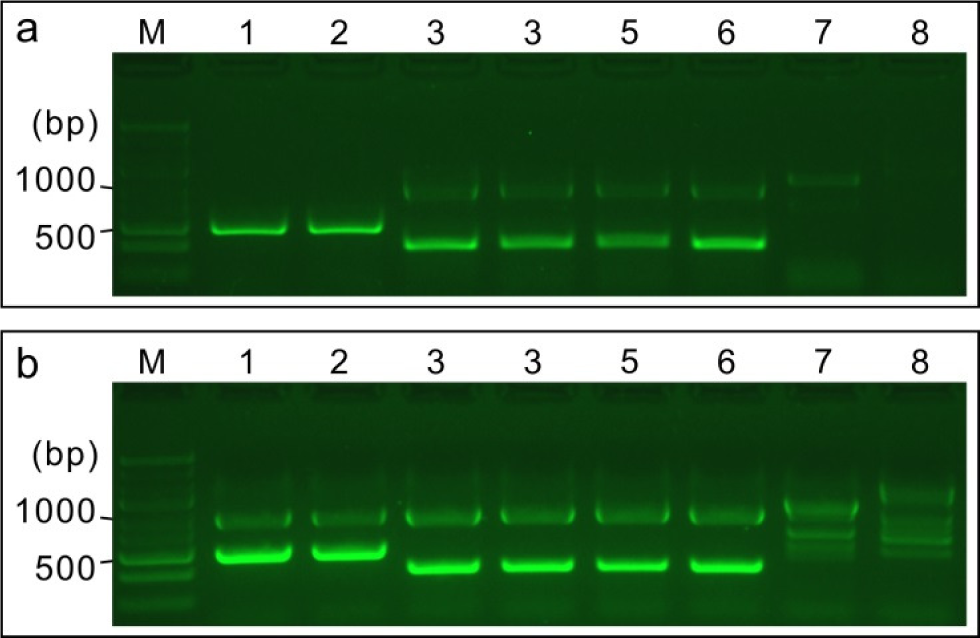
Analysis of selected fungal strains and canola samples by a multiplex PCR system developed by Inderbitzin et al. (2013). The reactions were conducted in Promega Go Taq Master Mix (a) or Thermo Scientific Phire Plant Direct PCR Master Mix (b). M, GeneRuler Express DNA Ladder. 1 and 2, Verticillium dahliae isolates APHL 232 and APHL 303, respectively. 3, V. longisporum isolate APHL 189. 4, The canola sample from which APHL 189 was isolated. 5, Canola sample 23CAN-305B. 6, Canola sample MB 2. 7, Canola sample 23CAN023. 8, Healthy canola.

## Discussion

### Specificity of P-VL and LAMP-VL

The specificity of P-VL and LAMP-VL are reflected by the specificity of their target genes, GenBank accession numbers CRK29573 and CRK26624, respectively. These two genes are present in *V. longisporum* only. Blasting the NCBI nr/nt database with the two genes did not produce any hit. When Blasting the two genes against the wgs database, eight hits with 100% query cover and 100% similarity were produced for both genes. These eight hits were from the eight whole-genome sequenced *V. longisporum* strains, respectively, and these eight strains represent all the whole-genome sequenced strains available in the database (Verified Jan 22, 2024). Thus, based on the sequence analysis, the two genes are specific to *V. longisporum* at the species level and ubiquitous at the strain level. This is in contrast with the other two *Verticillium*-specific genes (CRK12486 and CRK45493) identified in our initial candidate selection, which are present only in the six whole-genome sequenced A1/D1 strains (VLB2, VL20, VL2, VL1, Vl43 and Vl145c), but not in the two A1/D3 strains (PD589 and Vl32). It is likely that CRK29573 and CRK26624 are derived from the A1 parent and CRK12486 and CRK45493 are derived from the D1 parent. The specificity of P-VL and LAMP-VL was further demonstrated by the test on non-*V. longisporum* species. No signal was produced from any reaction.

### Sensitivity of P-VL and LAMP-VL

As a diploid, *V. longisporum* carries two copies of most genes, with each copy from one parent (Tran et al. 2013). The exception is the nuclear ribosomal region (rDNA), for which only one type is present in each lineage, as a result of DNA loss and/or concerted evolution (Depotter et al. 2016). For some genes, the two copies may have similar sequences that can be detected by the same set of primers. On the other hand, for many other genes, the two copies may have significant differences. This would result in a primer set for one copy unable to detect the other copy, but also Blast search of one copy would fail to identify the other one. In addition, there may be other genes that are present in only one parent and no homolog in the other parent. These genes are the real single-copied genes. We do not know whether CRK29573 and CRK26624 are real single-copied genes or genes with their homologs having different sequences. Nevertheless, based on the Blast results, these two genes are the single targets of P-VL and LAMP-VL, respectively, in the *V. longisporum* genome. Thus, the sensitivity of P-VL and LAMP-VL was expected to be lower than primers that target multiple-copied genes or DNA regions.

In the sensitivity test, the low limits for P-VL and LAMP-VL in a positive reaction were 28 copies of genomic DNA. Given the fact that the efficiency of DNA extraction from mycelia was not 100%, the low limit of the fungal cells for a positive reaction would be larger than 28. According to Yang et al. (2021), the extraction efficiencies of pathogen DNA from plant tissue was approximately 50%. Thus, we expected that the P-VL and LAMP-VL could detect *V. longisporum* DNA from as less as 56 cells. With the consideration that one DNA extraction generally results in 50 µ L of DNA and 2 µ L is used in one qPCR or LAMP reaction, the low limit of *V. longisporum* cell numbers in a sample from which the pathogen can be detected by our qPCR or LAMP assay is 1,400.

### The usefulness of the triplex qPCR and the LAMP

Two typical Verticillium stripe symptoms are dark unilateral stripes on the stems and microsclerotia in the stem cortex (Depotter et al. 2016). We have never observed these two symptoms on a single stem sample collected in our surveys which sampled randomly across Alberta and represented approximately 1% of canola acres in each County or Municipal District (n = approximately 380 fields, annually). In 2021-2023, canola stem samples showing symptoms suspicious for Verticillium stripe, such as discoloration on the internal stem tissues in a ‘starburst’ pattern, rather than sectoring, or unilateral striping on stems, were observed and tested for the presence of *V. longisporum*. However, the results were always negative (Harding et al. 2021; 2022; 2023). The results presented herein are the first confirmation of *V. longisporum* in our random Provincial survey. However, even on the sample 23CAN-305B from which *V. longisporum* was detected, the symptoms were not different from the symptoms of other samples from which only one or both of the two blackleg pathogens were detected. This was largely because this sample was infected by not only *V. longisporum* but also the two blackleg pathogens (Table 3). On the other hand, this also reflected the difficulty of symptom-based diagnosis of Verticillium stripe, at least on Alberta samples. Therefore, the triplex qPCR will greatly aid in *V. longisporum* detection from canola samples in Alberta, even when the two blackleg pathogens are not the detection targets. In our canola survey conducted in 2023, we tested 306 canola stem samples (including the twenty 23CAN samples in Table 2) with the triplex qPCR. All these samples had similar symptoms as the sample 23CAN-305B. *Verticillium longisporum* was detected from one sample only (23CAN-305B). In contrast, *L. biglobosa* or *L. maculans* was detected from 235 samples. These data again emphasized the difficulty in detecting *V. longisporum* visually when blackleg symptoms are present and the importance of using the triplex qPCR to test canola samples with ambiguous symptoms to discriminate between blackleg and Verticillium stripe.

The LAMP assay developed in this study can be used to confirm the results from the qPCR. Our data indicated that the LAMP had a similar sensitivity as the qPCR. Compared to qPCR, LAMP was believed to be more specific because LAMP targets six DNA regions by the six primers (Notomi et al. 2000). LAMP also has the advantage of being simple to conduct and no requirement for special equipment. Thus, LAMP was expected to be useful for on-site diagnosis. However, successful LAMP reactions are indeed highly dependent on the quality of template DNA, which has been overlooked by many researchers. DNA extracted by many simplified protocols (i.e. meant to be conducted in the field) employing high concentrations of detergents or alkalinity, should be avoided when producing template DNA for LAMP assays. Therefore, at the current time and for diagnostic labs equipped with qPCR cyclers, qPCR assays should be considered preferable to LAMP. This recommendation is further highlighted in the case of *V. longisporum* detection because successful extraction of good quality DNA from canola stems by simplified methods is more difficult than from other plant tissues.

In summary, in this study, we developed a triplex qPCR system that can be used to detect *V. longisporum*, *L. biglobosa,* and *L. maculans* in a single reaction. We confirmed the usefulness of this system on canola samples collected from different locations in Alberta. In addition, a LAMP assay was also developed, which can be used to verify the results from the triplex qPCR. We recommend the use of this triplex qPCR/LAMP combo system to others interested in detecting *V. longisporum* from plant samples.

